# Larger bacterial populations evolve heavier fitness trade-offs and undergo greater ecological specialization

**DOI:** 10.1101/2020.02.02.930883

**Authors:** Yashraj Chavhan, Sarthak Malusare, Sutirth Dey

## Abstract

Evolutionary studies over the last several decades have invoked fitness trade-offs to explain why species prefer some environments to others. However, the effects of population size on trade-offs and ecological specialization remain largely unknown. To complicate matters, trade-offs themselves have been visualized in multiple ways in the literature. Thus, it is not clear how population size can affect the various aspects of trade-offs. To address these issues, we conducted experimental evolution with *Escherichia coli* populations of two different sizes in two nutritionally limited environments and studied fitness trade-offs from three different perspectives. We found that larger populations evolved greater fitness trade-offs, regardless of how trade-offs are conceptualized. Moreover, although larger populations adapted more to their selection conditions, they also became more maladapted to other environments, ultimately paying heavier costs of adaptation. To enhance the generalizability of our results, we further investigated the evolution of ecological specialization across six different environmental pairs and found that larger populations specialized more frequently and evolved consistently steeper reaction norms of fitness. This is the first study to demonstrate a relationship between population size and fitness trade-offs and the results are important in understanding the population genetics of ecological specialization and vulnerability to environmental changes.

## Introduction

Adaptation of a biological population to a given environment may not concomitantly increase its fitness in other environments (Kassen, 2002; Anderson *et al.*, 2013; Kassen, 2014; Cooper, 2014; Bono *et al.*, 2017). In many cases, adaptation to one environment can also lead to maladaptation to others (Cooper and Lenski, 2000; Kassen, 2002; Bataillon *et al.*, 2011; Remold, 2012). Such incongruity in fitness changes across environments forms the basis of ecological specialization, which happens when the fittest type in one environment cannot be the fittest type in another environment (Fry, 1996), and tends to restrict the niche breadth of populations to a narrow range of environments (Levins, 1962, 1968; Futuyma and Moreno, 1988; Agrawal *et al.*, 2010). Studies of ecological specialization routinely invoke trade-offs in fitness across environments (Levins, 1962), with the latter leading to the intuitive and widespread assumption that the jack-of-all-trades is a master-of-none (MacArthur, 1984). Such fitness trade-offs and ecological specialization underlie the evolution of a wide range of biological properties and processes, including (but not restricted to) the composition of ecological communities (Kneitel and Chase, 2004; Farahpour *et al.*, 2018), host specificity in several systems (Rausher, 1984; Joshi and Thompson, 1995; Messina and Durham, 2015), virulence (Messenger *et al.*, 1999), resistance to a variety of agents like herbivores (Koricheva, 2002), parasites (Boots, 2011), antibiotics (Andersson and Hughes, 2010; MacLean *et al.*, 2010), etc.

The term ‘trade-off’ has been used in a variety of contexts in evolutionary studies (Agrawal *et al.*, 2010), with the following three main usages: (1) when a trait that is adaptive in a given environment is costly (maladaptive or detrimental to fitness) in others (Bell and Reboud, 1997; Kassen, 2014; Rodríguez-Verdugo *et al.*, 2014; Sane *et al.*, 2018); (2) unequal adaptation to alternative environments, wherein populations become adapted to all the environments under consideration but no single genotype can be the fittest one in all the environments (Bell and Reboud, 1997; Remold, 2012; Kassen, 2014); (3) life-history trade-offs across traits that arise within a single environment due to the systemic properties and constraints of organismal features (Stearns, 1989; Prasad *et al.*, 2001; Knops *et al.*, 2007). The first two of the above usages pertain to trade-offs in fitness across environments, which can directly give rise to ecological specialization. Although the physiological and molecular mechanisms of trade-offs and specialization are difficult to decipher in multicellular organisms (Agrawal *et al.*, 2010), there is a fairly detailed understanding of such mechanisms in microbes (Ferenci, 2016). However, the role of a key parameter like population size in determining fitness trade-offs, and the resultant specialization across a pair of environments, remains relatively unclear, even in asexual microbes.

Population size is known to shape a large variety of evolutionary phenomena and properties, including rate and extent of adaptation (Desai and Fisher, 2007; Desai *et al.*, 2007; Sniegowski and Gerrish, 2010; Chavhan *et al.*, 2019a), repeatability of adaptation (Szendro *et al.*, 2013; Lachapelle *et al.*, 2015), biological complexity (LaBar and Adami, 2016), efficiency of natural selection (Ohta, 1992; Petit and Barbadilla, 2009; Chavhan *et al.*, 2019a), etc. Numerous theoretical and empirical results have *indirectly* linked population size with the extent of ecological specialization. For example, multiple theoretical and empirical studies have established that larger populations generally adapt faster (Gerrish and Lenski, 1998; Desai and Fisher, 2007; Desai *et al.*, 2007; Sniegowski and Gerrish, 2010). Moreover, larger populations are expected to adapt primarily via rare large effect beneficial mutations while relatively smaller populations adapt slower through common beneficial mutations of modest effect sizes (reviewed in (Sniegowski and Gerrish, 2010)). Interestingly, the link between the size of a beneficial mutation and its deleterious pleiotropic effects has been touched upon by a rich array of theoretical studies. In Fisher’s geometrical model of adaptation on a multidimensional phenotype space where each orthogonal dimension represents a trait, larger mutational effect sizes are associated with greater probabilities of affecting multiple traits deleteriously (Fisher, 1930). This was the basis of Fisher’s argument for micromutationism, the idea that adaptation is largely driven by mutations of very small effects. Although Fisher’s original model assumed that all traits are equally important to fitness, it is easy to imagine cases where a few focal trait(s) affect(s) fitness while others do not (Orr and Coyne, 1992). In such a case, beneficial mutations with larger effects on the focal trait(s) would not only be selectively favoured, but also likely to be associated with large pleiotropic disadvantages to other traits. Along these lines, several theoretical studies assume that larger mutational benefits also have heavier pleiotropic disadvantages (Lande, 1983; Orr and Coyne, 1992; Otto, 2004). When we combine these two insights, a new testable hypothesis emerges: larger asexual populations should show greater specialization by adapting more specifically to their environment of selection and should also suffer heavier costs in alternative environments. To the best of our knowledge, there are no direct experimental tests of the relationships of such specializations and their underlying trade-offs with population size. To begin with, as pointed out repeatedly in the literature, experimental evolution studies of fitness trade-offs and the resulting specialization have not been conducted at variable population sizes (Kawecki *et al.*, 2012; (Bataillon, Thomas *et al.*, 2013; Cooper, 2014; Kraemer and Boynton, 2017). Furthermore, several recent evolution experiments with microbes which have provided important insights in this regard have focused on the pleiotropic profiles of individual mutations (reviewed in (Bono *et al.*, 2017)), and not on population-level properties like population size. To address this lacuna, here we use experimental evolution with *Escherichia coli* populations of different sizes to test if larger populations evolve bigger fitness trade-offs and specialize more across environments.

To avoid semantic ambiguity, we follow Fry (1996) and define specialization across two environments as any case where the reaction norms for fitness intersect with each other. Evolutionary experiments have conventionally studied fitness trade-offs across environments from three major perspectives: (1) as negative correlations in fitness of populations selected in two or more distinct environments (Bell and Reboud, 1997; Jessup and Bohannan, 2008; Lee *et al.*, 2009); (2) as fitness deficits below the ancestral levels (costs of adaptation) in the alternative environment(s) that accompany adaptation to the environment in which evolution takes place (Lee *et al.*, 2009; Bono *et al.*, 2017); (3) as differences in fitness across environmental pairs (Kassen, 2014; Schick *et al.*, 2015). Although these perspectives can potentially be related, they are clearly not equivalent, and therefore might lead to different insights about the process of ecological specialization. For example, ecological specialization can happen with or without costs of adaptation (see Appendix S1 and Fig. S1 in Supplementary Information for details). Therefore, we decided to use all above three perspectives to investigate how population size affects fitness trade-offs and the resulting specialization across environments. To this end, we propagated replicate *E. coli* populations of two different sizes in two different nutritionally limiting environments (Galactose minimal medium and Thymidine minimal medium). We note that these experiments can also be contextualized in terms of a combination of two other questions: (A) Do pleiotropic effects act reciprocally across the pair of environments in question? (B) Are pleiotropic effects correlated to the corresponding direct effects on fitness? Although several previous studies have investigated these questions separately, they have not combined them in order to look at the effects of population size on fitness trade-offs. With regards to the reciprocity of trade-offs, for example, collateral sensitivity profiles have been shown to be reciprocal across several pairs of antibiotics (Imamovic and Sommer, 2013). Contrastingly, other studies have found asymmetric (and not reciprocal) trade-offs across pairs of environments (Travisano, 1997; Lee *et al.*, 2009). Along similar lines, previous studies have reported evidence of both high and low correlations of pleiotropic and direct fitness effects (Ostrowski *et al.*, 2005; Sane *et al.*, 2018). Linking the above two questions, we also sought to determine if our experimental populations evolved reciprocal trade-offs and whether direct fitness gains in one environment led to correlated disadvantages in the other.

We found that, descending from a common ancestor, our experimental populations evolved a strong negative correlation between fitness across the two environments. As expected, the larger populations adapted more to the selection environments. Interestingly, the larger populations also paid heavier costs of adaptation. We further assayed the fitness of the evolved populations in two more nutritionally limiting environments, which enabled us to quantify the extent of specialization across six environmental pairs. Remarkably, we found that the larger populations specialized more, evolving steeper reaction norms of fitness. To the best of our knowledge, this is the first study to directly test the effects of population size on ecological specialization brought about by fitness trade-offs.

## Materials and Methods

### Experimental Evolution

We founded 24 populations from a single *E. coli* MG1655 colony and propagated them for ~ 480 generations at two different population sizes: Large (L) or Small (S) (defined below). For each population size, we had two different kinds of environments: Thymidine (T) or Galactose as the sole carbon source in a M9-based minimal medium (for details, see Appendix S2 in Supporting Information). This 2 × 2 design gave rise to four population types (TL, TS, GL, and GS) where the first letter represented the only carbon source in the selection environment, and the second letter represented the population size. We chose these carbon sources because they have very different metabolic pathways in *E. coli* (Frey, 1996; Loh *et al.*, 2006; Díaz-Mejía *et al.*, 2009; Barupal *et al.*, 2013). Furthermore, galactose and thymidine differ in terms of their uptake within *E. coli* cells. Specifically, whereas the uptake of thymidine occurs via NupG and NupC proteins (Patching *et al.*, 2005), galactose uptake is mediated by GalP (Henderson *et al.*, 1992). Taken together, based on disparate metabolism and uptake mechanisms of the two carbon sources, we expected *E. coli* populations to show fitness trade-offs across them. Each population type had six independently evolving replicate populations. We propagated all the 24 populations using the standard batch-culture technique at a volume of 300 μl in 96 well plates shaking continuously at 150 rpm at 37° C. The large populations (TL and GL) had a periodic bottleneck of 1:10 while and the small ones (TS and GS) faced a periodic bottleneck of 1:10^4^. To ensure that our treatments did not spend vastly different amounts of time in the stationary phase, we bottlenecked the large populations every 12 hrs (every 3.3 generations), and the small ones every 48 hrs (every 13.3 generations). Overall, L and S corresponded approximately to 9.9 × 10^7^ and 3.9 × 10^5^ respectively in terms of the harmonic mean population size (Lenski *et al.*, 1991). In terms of the measure of population size relevant for the extent of adaptation in asexual populations (Chavhan *et al.*, 2019a), L and S corresponded approximately to 9.0 × 10^6^ and 2.2 × 10^3^ respectively.

### Quantification of fitness and trade-offs across environments

At the end of our evolution experiment, we revived the cryostocks belonging to each experimental population in glucose-based M9 minimal medium and allowed them to grow for 24 hours. Next, we performed automated growth-assays on each of the 24 revived populations using a well-plate reader in multiple environments (Synergy HT, BIOTEK ® Winooski, VT, USA). Using optical density at 600 nm as the measure of population density, we obtained growth readings every 20 minutes for 24 hours. We ensured that the physical conditions during the assays were identical to culture conditions (96 well plates shaking at 150 rpm at 37° C). Using a randomized complete block design (RCBD), we conducted the fitness measurements over six different days, assaying one replicate population of each type in both the environments on a given day (Milliken and Johnson, 2009). We estimated fitness as the maximum growth rate (R) (Kassen, 2014; Ketola and Saarinen, 2015; Vogwill *et al.*, 2016), which was computed as the maximum slope of the growth curve over a moving window of ten readings (Leiby and Marx, 2014; Karve *et al.*, 2015, 2016, 2018; Chavhan *et al.*, 2019a; Chavhan *et al.*, 2019b).

We labelled the environment in which selection occurred as ‘home’ and the other (alternative) environment(s) as ‘away.’ The presence of the common ancestor as the reference against which fitness gains or reductions could be tested allowed us to differentiate between ecological specialization and costs of adaptation. We studied trade-offs and ecological specialization from three major conventional perspectives:

1. If fitness trade-offs exist between two environments, selection is expected to result in strong negative correlations in fitness across them (Kassen, 2014). Therefore, we determined if relative fitness in Galactose (henceforth “Gal”) had a significant negative correlation with relative fitness in Thymidine (henceforth “Thy”).
2. We also determined if our experimental populations paid significant costs of adaptation. To this end, we first established whether our experimental populations had adapted significantly to their home environment (Thy for TL/TS and Gal for GL/GS). Next, we determined if the populations had maladapted significantly to their away environment (Gal for TL/TS and Thy for GL/GS). For this, we normalized all fitness values in a given environment by the ancestral fitness in the corresponding environment, which is equivalent to scaling the ancestral fitness value to 1 (Kassen, 2014). We then used single sample t-tests to ascertain if the fitness of a given population type differed significantly from the ancestor. We corrected for inflations in family wise error rate using the Holm-Šidák procedure (Abdi, 2010). We also computed Cohen’s *d* to analyse the statistical significance of these differences in terms of effect sizes (Cohen, 1988). We concluded that a population had paid a cost of adaptation only if it had adapted significantly to its home environment (i.e., scaled fitness > 1) and simultaneously maladapted significantly to its away environment (i.e., scaled fitness < 1). We used a mixed model ANOVA with RCBD to analyse if population size and the identity of the home environment interacted with each other statistically to shape the fitness in home environment (henceforth Fitness_home_). To this end, we used ‘Population Size’ (two levels: L or S) and ‘Home environment’ (two levels: Gal or Thy) as fixed factors crossed with each other, and ‘Day of assay’ (six levels: 1 to 6) as the random factor. We also analysed the effect size of the main effects using partial η^2^, interpreting the latter as representing small, medium, or large effect for Partial η^2^ < 0.06, 0.06 < Partial η^2^ < 0.14, 0.14 < Partial η^2^ respectively (Cohen, 1988). Furthermore, we analyzed if the relative extent of fitness loss in the away environment (= 1 - Fitness_away_) was significantly different for the large and small populations. To this end, we used a mixed-model ANOVA (RCBD) with ‘Population Size’ (two levels: L or S) and ‘Home-Away pair’ (two levels: Gal-Thy or Thy-Gal) as fixed factors crossed with each other, and ‘Day of assay’ (six levels: 1 to 6) as the random factor.
3. We quantified the environmental specificity of adaptation using differences in the relative fitness of experimental populations across different home-away environmental pairs. This quantity represents the difference in the degrees to which a population adapts to the two environments under consideration (Remold, 2012). Such a difference between home- and away- relative fitness values (= Fitness_home_ – Fitness_away_) can be represented graphically as slopes of reaction norms of fitness. Since the quantification of reaction norm slopes does not require selection in each assay environment, we enhanced the generalizability of our study by assaying the fitness of all the 24 evolved populations (TL, TS, GL, and GS) in two more nutritionally limited environments (Maltose minimal medium (henceforth “Mal”) and Sorbitol minimal medium (henceforth “Sor”)). This allowed us to compare the reaction norm slopes of large and small populations across six home-away environmental pairs (Thy-Gal, Thy-Mal, Thy-Sor for TL and TS; Gal-Thy, Gal-Mal, Gal-Sor for GL ad GS).

We first determined if a population type had specialized significantly across a given home-away pair. We followed Fry (1996) to identify specialization across a pair of environments as any case where the population’s reaction norm intersected with the corresponding ancestral reaction norm. This would happen whenever the unambiguous fittest type in one environment is not unambiguously the fittest type in the other environment (Fig. 1). To identify cases of specialization, we first determined if the population type in question had significantly different Fitness_home_ relative to the common ancestor. Next, we determined if the population type’s Fitness_away_ was significantly different from that of the ancestor. If the population type increased its fitness significantly in both its home and away environments, its reaction norm would not intersect with the ancestral norm. This would imply lack of ecological specialization. On the other hand, if the population type’s fitness increased significantly in its home environment but failed to do so in its away environment, its reaction norm would intersect with the ancestral norm, revealing significant specialization across the environmental pair in question (Fig. 1). We performed this procedure for each of the four population types to determine if specialization had occurred across the six home-away pairs under consideration.

We also compared the reaction norm slopes of each of the four population types with that of the ancestor across all the home-away pairs under consideration. To this end, we conducted single sample t-tests against the ancestral level (ancestral reaction norms have zero slope), followed by correction for family-wise error rates using the Holm-Šidák procedure.

To further study how population size affects the specificity of adaptation, we determined if the reaction norm slopes were significantly different across large and small populations. To this end, we conducted two separate mixed model ANOVAs (RCBD), one for selection in Thy and the other for selection in Gal. The design of these mixed model ANOVAs had ‘Population Size’ (two levels: L or S) and ‘Home-Away pair’ (three home-away pairs) as fixed factors crossed with each other, and ‘Day of assay’ (six levels: 1 to 6) as the random factor.

**Fig. 1.**
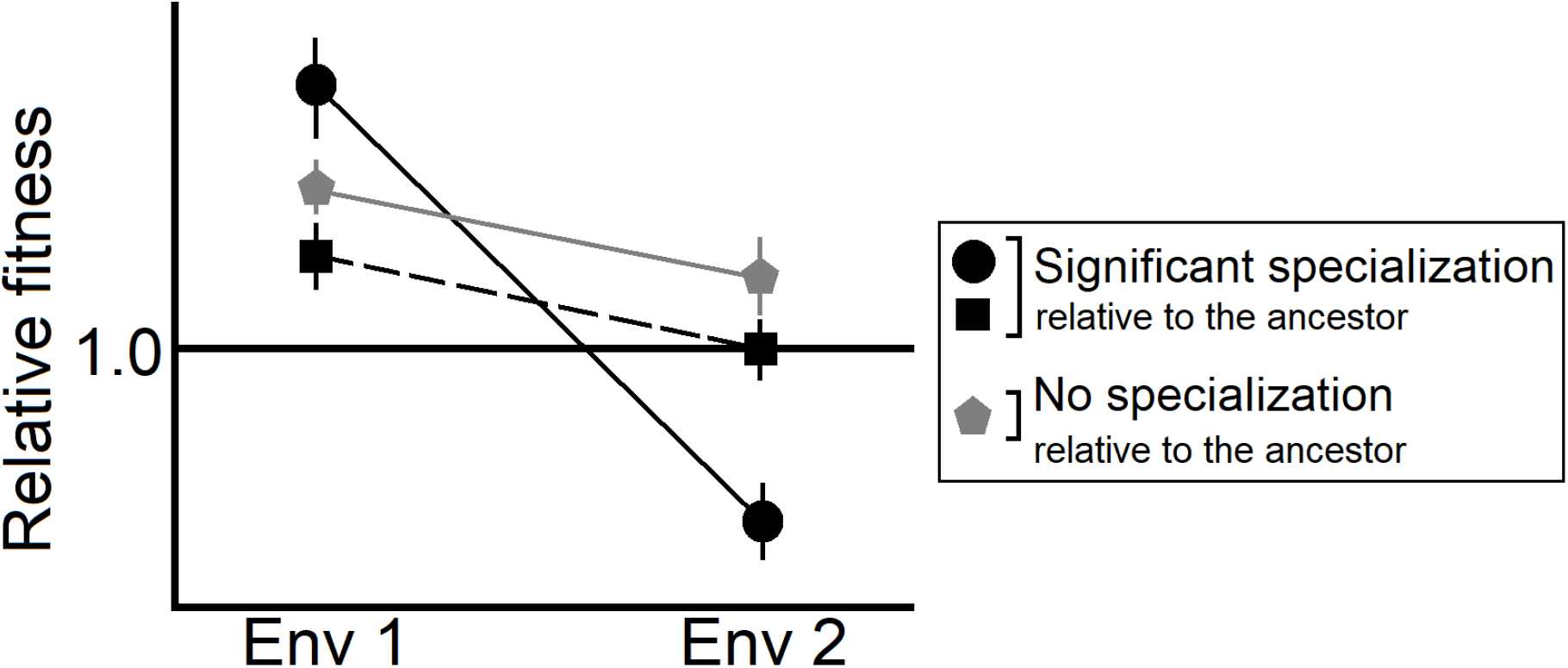
Schematic representation of ecological specialization across two environments. All the fitness values are scaled by the ancestral value (=1 (horizontal black line)). The error bars represent 95% confidence intervals. Significant ecological specialization occurs when the reaction norms of fitness intersect (which happens when the unambiguous fittest type in one environment is not the unambiguous fittest type in the other environment).

## Results

### 1. Fitness trade-offs as negative correlations: Fitness in Gal was negatively correlated with fitness in Thy

We found a strong negative correlation between fitness in Gal and fitness in Thy (Fig. 2; Spearman’s ρ = −0.744; *P* = 3.04 × 10^−5^). This negative correlation revealed that the fittest type in Gal was never the fittest type in Thy (compare Fig. 2 with Fig. S1), which implied the occurrence of ecological specialization in our experimental populations.

**Fig. 2.**
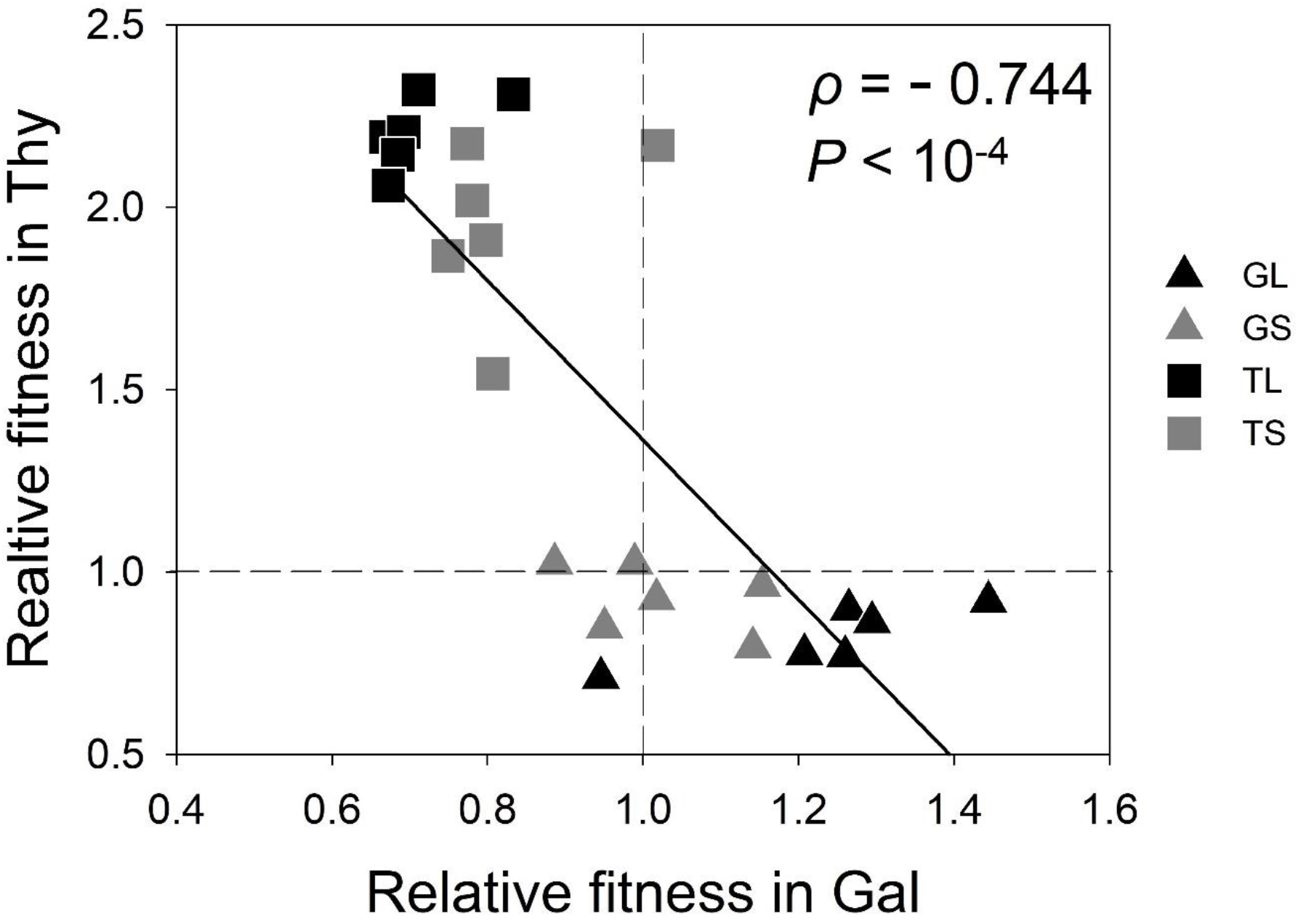
Correlation between relative fitness values in Galactose and Thymidine minimal media after evolution in these environments at two different population sizes. The black line represents the best linear fit (R² = 0.63); the dotted lines represent ancestral levels of fitness.

Since trade-offs based on negative fitness correlations can potentially happen with or without costs of adaptation (Fig. S1; Fry (1996); Kassen (2014)), we next determined if our populations had evolved significant costs of adaptation. We also examined whether larger populations had evolved greater costs of adaptation.

### 2. Trade-offs as costs of adaptation: Larger populations paid more costs of adaptation

Following Kassen (2014), we defined costs of adaptation as the simultaneous occurrence of fitness increase (adaptation) in the home environment and fitness decrease in the away environment (maladaptation). A comparison of Fig. 2 with Fig. S1 shows that the negative correlation between relative fitness values in Gal and Thy was accompanied by costs of adaptation.

To begin with, we found a significant main effect of population size on adaptation to the home environment, with the larger populations showing higher Fitness_home_ than the smaller ones (mixed model ANOVA: population size (F_1,15_ = 10.998, *P* = 0.005, η^2^ = 0.423 (large effect)). We also found a significant main effect of the home environment (F_1,15_ = 176.969, *P* = 1.044 ×10^−9^, η^2^ = 0.921 (large effect)). Importantly, we did not find a significant population size × home environment interaction (F_1,15_ = 0.102, *P* = 0.753).

We found that evolution in Thy resulted in significant costs of adaptation in case of both large (TL) and small (TS) populations (Table 1), i.e., both TL and TS adapted to Thy but became maladapted to Gal (Fig. 2; Table 1). However, evolution in Gal incurred significant costs of adaptation only in case of the large populations (GL); i.e., the GL populations adapted to Gal but became maladapted to Thy (Fig. 2; Table 1). The small populations that evolved in Gal (GS) neither adapted significantly to Gal nor became significantly maladapted to Thy (Fig. 2; Table 1). Thus, the GS populations did not pay any costs of adaptation.

**Table 1.** Occurrence of adaptation and maladaptation events in population types selected in Gal and Thy separately. The Holm-Sidak corrected *P* values correspond to single sample t-tests against the ancestral levels of fitness (= 1). *P* < 0.05 are shown in boldface. All the cases with *P* < 0.05 were also found to have large effect sizes (See Table S1).

| Population type | Fitness change in Gal | Fitness change in Thy | Cost of adaptation |
| --- | --- | --- | --- |
| TL | Maladaptation<br>**$P = 2.683 \times 10^{-4}$** | Adaptation<br>**$P = 3.240 \times 10^{-6}$** | Yes |
| TS | Maladaptation<br>**$P = 0.007$** | Adaptation<br>**$P = 7.327 \times 10^{-4}$** | Yes |
| GL | Adaptation<br>**$P = 0.049$** | Maladaptation<br>**$P = 0.013$** | Yes |
| GS | Not significant<br>$P = 0.617$ | Not significant<br>$P = 0.122$ | None |

Hence, larger populations paid significant costs of adaptation in both the environments, but the smaller ones did so only in one of the two environments under consideration.

Next, we determined if larger populations also had a higher magnitude of loss in relative fitness below the ancestral levels in their away environments. Indeed, we found that the larger populations lost significantly greater fitness than the smaller ones in their away environments (Fig. 3, mixed-model ANOVA: population size (main effect) F_1,15_ = 9.558, *P* = 0.007, partial η^2^ = 0.389 (large effect); home environment (main effect) F_1,15_ = 9.650; *P* = 0.007 partial η^2^ = 0.391 (large effect), population size × home environment (interaction) F_1,15_ = 0.002, *P* = 0.963). This result also implies that the effect of population size on Fitness_away_ would be the opposite of its effects on Fitness_home_. Table 1 also shows that adaptation in Thy (in case of TL and TS) was always accompanied by maladaptation in Gal; analogously, populations that adapted significantly to Gal showed significant maladaptation in Thy. This demonstrates that the fitness trade-offs were reciprocal across the Thy-Gal environmental pair.

**Fig. 3.**
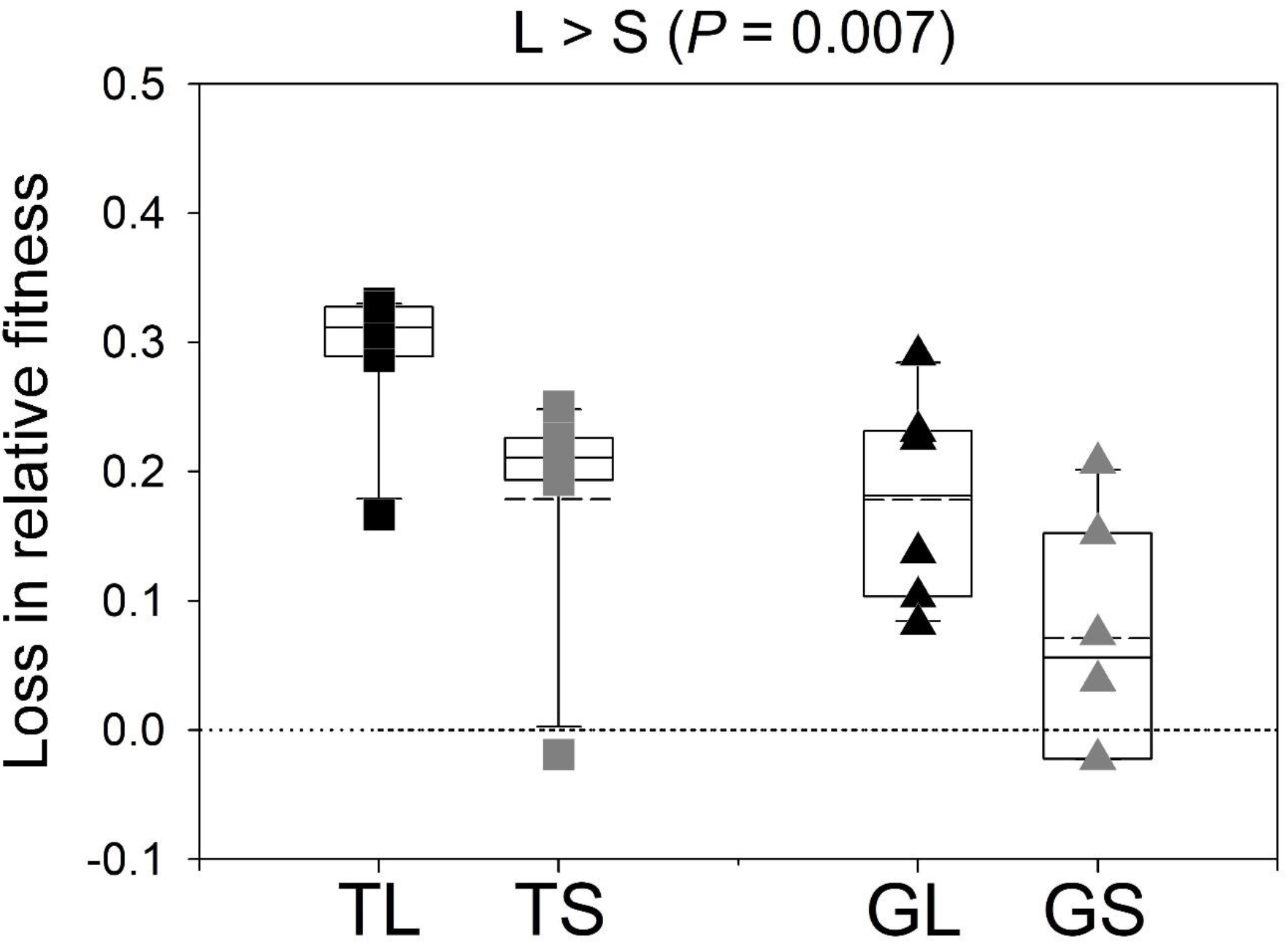
Loss of fitness below the ancestral levels in the away environments. In each case, the loss in relative fitness was computed as the difference between the descendant population’s relative fitness and the ancestor’s relative fitness. L and S represent large and small populations, respectively. The solid lines in the box plots mark the 25^th^, 50^th^, and 75^th^ percentiles while the whiskers mark the 10^th^ and 90^th^ percentiles; the dashed lines represent means (N = 6). The dotted line represents no loss from the ancestral fitness levels. See the text for details.

Overall, larger population adapted more to their home environments while paying greater costs of adaptation. This made them significantly more maladapted to the away environments.

Thus, as compared to the small populations, the large populations lost more Fitness_away_. We note that the magnitudes of such fitness decline cannot reflect the full extent of ecological specialization. This is because the former only represent deficits in Fitness_away_ whereas ecological specialization involves a difference between the extents of adaptation across the home and away environments. Thus, as the next logical step, we set out to measure the extent of specialization in our experimental populations.

### 3. Fitness trade-offs as extents of ecological specialization: Larger populations specialized more

We used reaction norm slope as a measure of the specificity of adaptation, computed as the difference between Fitness_home_ and Fitness_away_. As described earlier, we assayed the fitness of our experimental populations in two more nutrient-limited environments, which allowed us to compute the reaction norm slope across six different environmental pairs (three for Gal-selected populations and three for Thy-selected populations). Moreover, the normalization of fitness in any given environment with the corresponding ancestral value ensured that the slope of the ancestral reaction norms of relative fitness is zero across all environmental pairs (Kassen, 2014).

First, we determined whether our experimental populations had specialized significantly to their home environment. Following a previous study (Fry, 1996), we identified the evolution of specialization as the intersection of the reaction norms of the descendant treatments with that of the ancestor (Fig. 1).

On the one hand, the TL and TS populations had significantly greater fitness than the ancestor in their home environment (Table 1). On the other hand, the TL and TS populations had lower fitness than the ancestor in all the three away environments under consideration (Table S1). This reveals that the average reaction norms of both TL and TS populations intersected with the ancestral norms across all the three environmental pairs under consideration (T-Gal, Thy-Mal, Thy-Sor) (Fig. 4a). Hence, both the TL and TS populations had specialized significantly across all the three home-away pairs.

**Fig. 4.**
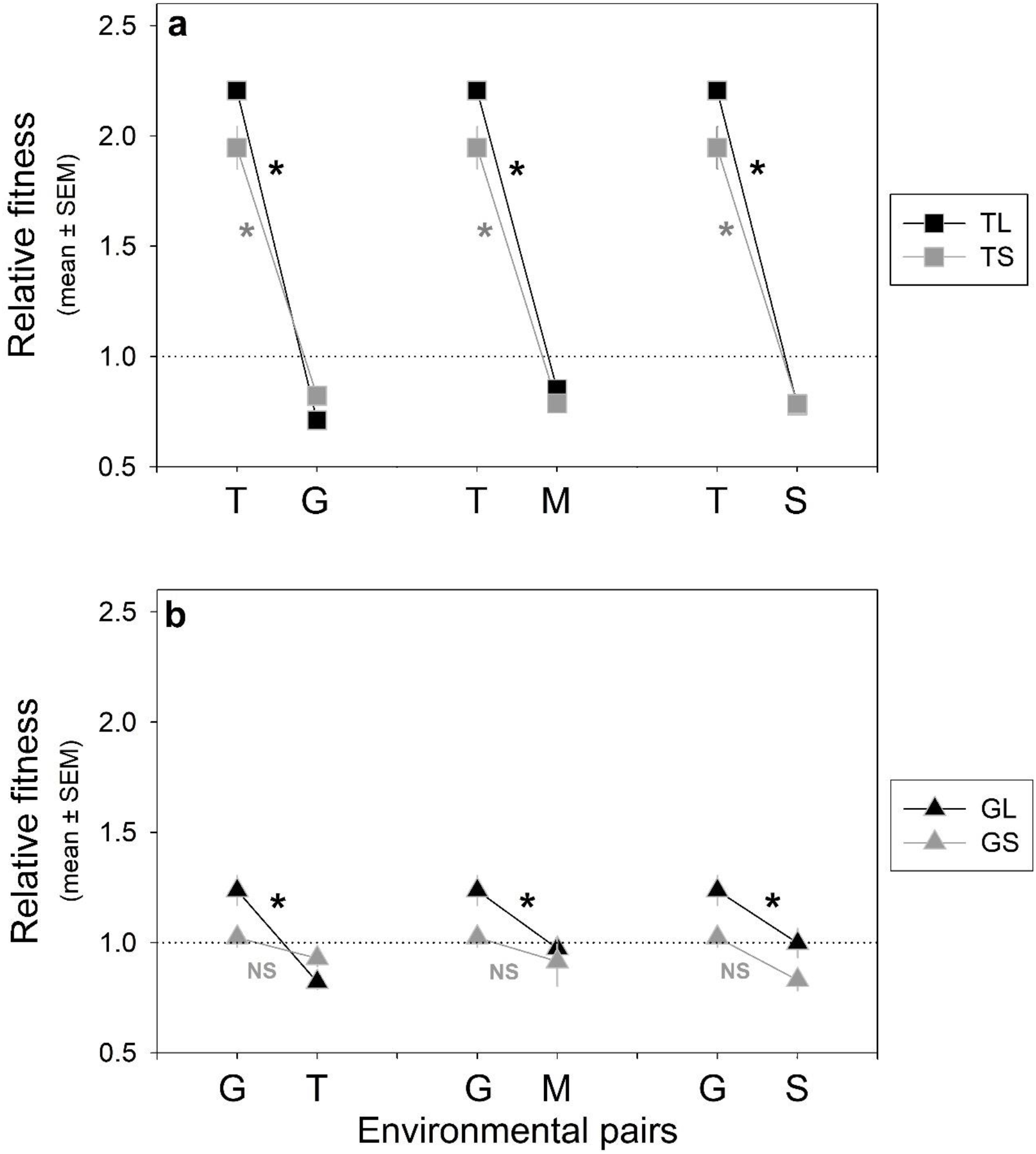
Reaction norms of fitness across the six home-away environmental pairs used in our study. The error bars represent SEM (N = 6). The asterisks represent significant ecological specialization with respect to the ancestor for the corresponding home-away pair; ‘NS’ denotes that the corresponding ecological specialization was not significant (see Table S1 and the text for details). The dotted lines represent ancestral reaction norms. Significant specialization happened when the reaction norms of a treatment population intersected the ancestral norm **(a)** Reaction norms for populations selected in Thy (TL (large) and TS (small). **(b)** Reaction norms for populations selected in Gal (GL (large) and GS (small). Also see Fig. 5 for reaction norm slopes across the six environmental pairs.

The large populations evolved in Gal (GL) had adapted significantly to their home environments (Table 1). Furthermore, the relative fitness of GL was not significantly greater than the ancestor in any of the three away environments (Table S1). Combining these pieces of information, GL populations specialized significantly (i.e., the fittest type in home environment was not the unambiguous fittest type in the away environment) across all the three home-away pairs (Gal-Thy, Gal-Mal, Gal-Sor) (Fig. 4b). Interestingly, the small populations evolved in Gal (GS) did not have significantly different fitness as compared to the ancestor in any of the four environments under consideration (Table S1). This implies that the GS populations did not specialize significantly across any of the three home-away pairs under consideration (Fig. 4b).

Amongst the Thy-selected populations, we found that both the large (TL) and small (TS) populations evolved significantly greater reaction norm slopes than that of the ancestor (i.e., reaction norm slope = 0) ((Fig. 5a, Table S2).

**Fig. 5.**
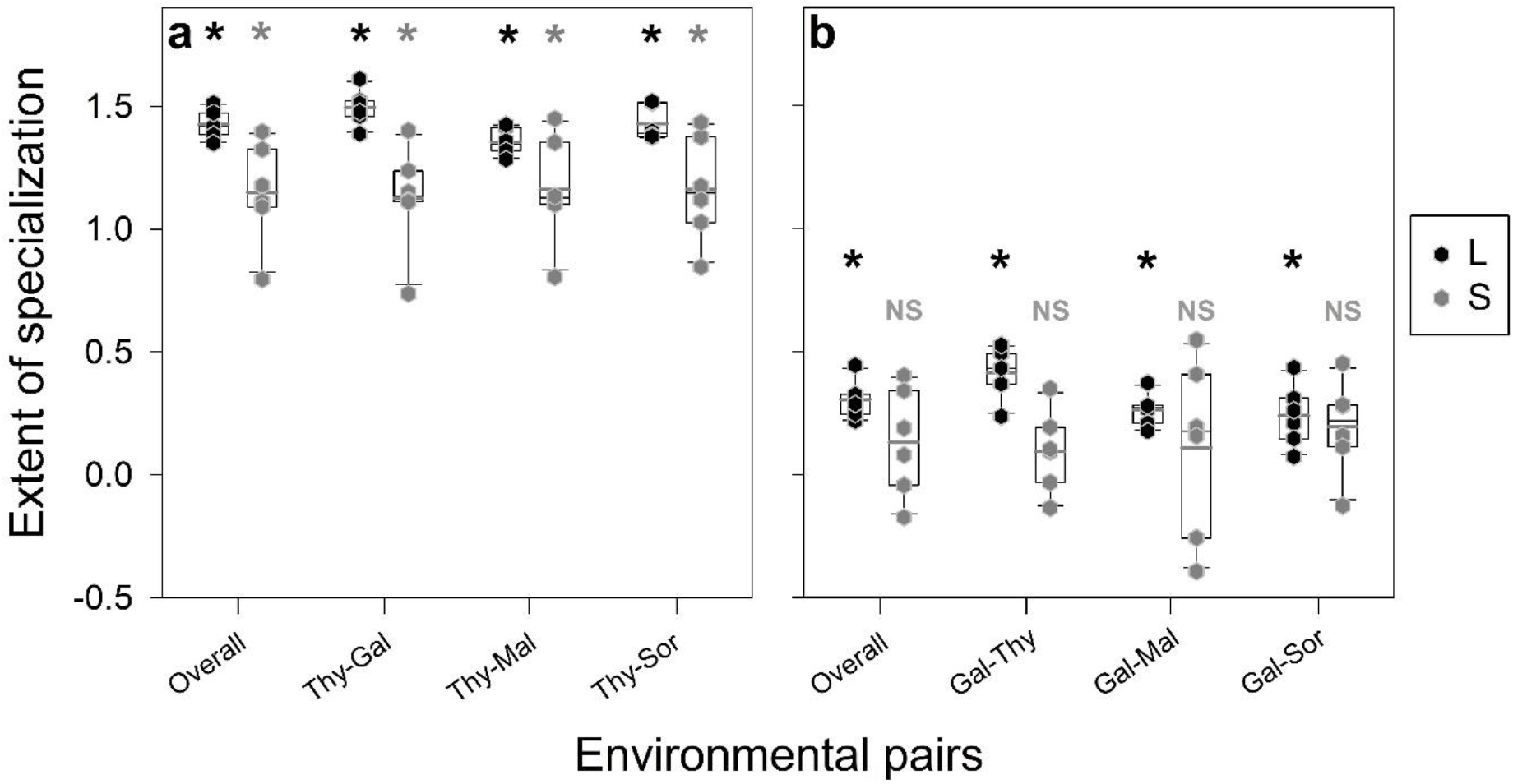
Slopes of reaction norms of the fitness of our experimental populations across six environmental pairs. The asterisks represent significant differences (single sample t-tests (*P* < 0.05)) from the ancestral slope (= 0); ‘NS’ denotes that the corresponding slopes are not significantly different from 0. The solid lines in the box plots mark the 25^th^, 50^th^, and 75^th^ percentiles while the whiskers mark the 10^th^ and 90^th^ percentiles; the thick horizontal lines represent means (N = 6). See the text and Table S2 for details and Fig. 4 for reaction norms across the six environmental pairs. **(a)** Populations evolved in Thy: reaction norm slopes (L > S (*P* < 10^−6^)). **(b)** Populations evolved in Gal: reaction norm slopes (L > S (*P* < 10^−2^)). Overall, the larger populations had steeper reaction norms.

Amongst the Gal-selected populations, we found that the GL populations had significantly steeper reaction norms than the ancestor across all the three home-away pairs under consideration (Fig. 5b; Table S2). However, the reaction norm slopes of the GS populations were not significantly different from that of the ancestor across any of the three home-away pairs (Fig. 5b; Table S2).

We further determined if larger populations had steeper reaction norms than smaller populations over the six different home-away environmental pairs. Indeed, we found a significant main effect of population size, with the larger populations evolving steeper reaction norms than the smaller ones; importantly, population size and home-away pair did not show significant statistical interaction (Fig. 5: mixed-model ANOVA for Thy-selected lines: Population Size (main effect) F_1,25_ = 43.664, *P* = 6.357 × 10^−7^, partial η^2^ = 0.636 (large effect); Home-Away pair (main effect) F_2,25_ = 0.566, *P* = 0.575; Population Size × Home-Away pair (interaction) F_2,25_ = 1.471, *P* = 0.249; mixed-model ANOVA for Gal-selected lines: Population Size (main effect) F_1,25_ = 8.147, *P* = 0.009, partial η^2^ = 0.246 (large effect); Home-Away pair (main effect) F_2,25_ = 0.428, *P* = 0.657; Population Size × Home-Away pair (interaction) F_2,25_ = 1.721, *P* = 0.199).

Overall, the large populations not only specialized more frequently (six home-away pairs for the large populations versus three for the small ones), they also evolved higher magnitudes of specificity of adaptation.

## Discussion

Fitness trade-offs across environments and the ensuing ecological specialization play key roles in understanding a variety of important phenomena, including the maintenance of biodiversity, local adaptation, etc. (reviewed in (Futuyma and Moreno, 1988; Agrawal *et al.*, 2010)). However, little is known about the relationship between fitness trade-offs and population size, even in relatively simple organisms like microbes.

In this study, we conducted experimental evolution to directly test how population size influences fitness trade-offs and the resulting ecological specialization. Inconsistencies in the usage of terms like trade-offs and costs of adaptation in the evolutionary biology literature complicate comparisons across studies. Therefore, here we investigated fitness trade-offs with three different (but not necessarily independent) perspectives. Our primary result is that regardless of how we chose to visualize trade-off, larger populations suffered more fitness trade-offs and thus evolved higher extents of ecological specialization. To the best of our knowledge, this is the first study to address and experimentally demonstrate this relationship between population size and fitness trade-offs.

The theory of adaptive dynamics in asexual populations predicts that while larger populations adapt primarily via rare large effect beneficial mutations, such mutations remain largely inaccessible to small populations, which adapt via mutations of relatively smaller effect sizes (Sniegowski and Gerrish, 2010; Chavhan *et al.*, 2019a). Moreover, a large body of studies suggests that mutational benefits of large sizes lead to heavier disadvantages due to antagonistic pleiotropy (Orr and Coyne, 1992; Otto, 2004; Griswold, 2007; Hague *et al.*, 2018). Our result that larger population pay higher costs of adaptation can thus be explained by a combination of the above two ideas. In other words, the notion that larger populations adapt primarily via large effect beneficial mutations which, in turn, are expected to lead to higher pleiotropic disadvantages, can potentially explain why larger population paid higher costs of adaptation which resulted in steeper reaction norms (Figs. 5 and S2). Furthermore, since multiple beneficial mutations can simultaneously rise to high frequencies in large populations (Desai and Fisher, 2007; Desai *et al.*, 2007), our observations can also be explained by the pleiotropic effects of a higher number of beneficial mutations in the larger experimental populations (TL and GL). Although our observations align with the underlying assumption that larger-effect beneficial mutations should be associated with heavier pleiotropic disadvantages, we have not presented direct genetic evidence for this assumption. We note that the small experimental populations in our study (TS and GS) harboured enough individuals to undergo clonal interference between distinct beneficial mutations while the larger populations (TL and GL) likely underwent adaptation based on multiple beneficial mutations within 480 generations (Desai and Fisher, 2007). Although the timescale of 480 generations is long enough to lead to substantial fitness increase, it is likely to be inadequate for the fixation of multiple beneficial mutations (Cooper, 2018). Thus, our experimental populations are expected to be genetically heterogeneous. Hence, any genetic test of the underlying assumption regarding the scaling of pleiotropy with the direct effects of the individual mutations in our experiment would need to account for the genetic background (epistatic interactions) and allelic frequencies at several loci. Although interesting in its own right, such an investigation was out of the scope of the current study.

A previous study with a collection of single beneficial mutations had found their pleiotropic effects in terms of carbon usage to be rarely antagonistic (Ostrowski *et al.*, 2005), which contrasts with our observations. Below we briefly speculate on the basis of these differences. Ostrowski *et al.*, (2005) had determined the pleiotropic effects of single beneficial mutations that had started rising in frequency during selection in glucose minimal medium (Rozen *et al.*, 2002). Glucose is known to be the preferred carbon source for *E. coli* (Brückner and Titgemeyer, 2002; Görke and Stülke, 2008). This preference for glucose is a likely outcome of metabolic optimization during the evolutionary history of these bacteria. Thus, the scope for adaptation on glucose should be relatively lower as compared to that on other carbon sources (like Thy and Gal). Since lower scope for adaptation is associated with slower increase in fitness (Couce and Tenaillon, 2015), adaptation in glucose minimal medium should be relatively slower. Indeed, it took more than 1000 generations for the populations in Lenski’s Long Term Evolution Experiment to increase their growth rate by ~30% on glucose as the sole carbon source (Novak *et al.*, 2006). Contrastingly, even the small populations of our study could increase their growth rate in Thy by >70% within 480 generations (this increase was >100% for the large populations). As compared to Thy, Gal is closer to glucose in terms of metabolic pathways, suggesting that the adaptation in Gal should be slower than that in Thy. Our observations agree with this expectation (~20% increase in GL and no significant increase in GS) (Table 1). Furthermore, we note that the average fitness effect of mutations enriched on glucose for 400 generations reported in Ostrowski *et al.*, (2005) was approximately 10%. Taken together, the direct effects of beneficial mutations that drove adaptation in our study (in TL, TS, and GL, but not in GS) are expected to be greater than those reported in Ostrowski *et al.*, (2005). The small size of direct fitness in their study could be a potential reason behind the lack of detectable antagonistic pleiotropic effects. Thus, the notion that larger direct effects are associated with heavier pleiotropic disadvantages could be enough to explain why our results are different from Ostrowski *et al.*, (2005). Lastly, since GL and TL populations are large enough to have multiple beneficial mutations within the same lineage rising simultaneously to high frequencies, the substantial pleiotropic costs in these populations are likely to be associated with multiple mutations as opposed to single mutations as reported in Ostrowski *et al.*, (2005).

We briefly note that although maladaptation to the away environments can potentially be caused by the accumulation (via drift in the home environment) of neutral mutations that are contextually deleterious in the away environments, it is unlikely to be an explanation of our observations. There are two key reasons behind this assertion. First, a period of 480 generations is too small for mutation accumulation to show its phenotypic effects in clonally derived bacterial populations with harmonic mean population sizes greater than 10^4^ (which happens to be the case here) (Kassen, 2002; Cooper, 2018). Second, the effects of accumulation of conditionally neutral mutations are not expected to be different across populations of different sizes (Kimura, 1983; Hall and Colegrave, 2008).

Our study shows that when the environment remains constant for long periods, adaptation in larger numbers can make populations more specialized to this environment. This relationship between population size and ecological specialization has many important implications. Foremost, owing to their higher extent of specialization, larger populations can become vulnerable to sudden changes in the environment, as predicted by a recent study (Chavhan *et al.*, 2019b). Interestingly, if the environment abruptly shifts between two states that show fitness trade-offs with each other, then populations with a history of evolution at larger numbers would be at a greater disadvantage than historically smaller populations. Microbial populations routinely experience such abrupt shifts across environmental states that are known to show fitness trade-offs with each other. For example, costs of antimicrobial resistance are expected to check the spread of resistant microbes if antimicrobials are removed abruptly from the environments (Andersson and Hughes, 2010; Hill *et al.*, 2015). Moreover, pathogens are also expected to experience fitness trade-offs when they migrate across different hosts (Turner and Elena, 2000; Smith-Tsurkan *et al.*, 2010).

Our results also predict that in the face of environmental changes, larger populations may not adapt better than smaller ones. Pleiotropy has been routinely invoked to explain why evolution should mostly proceed via small effect mutations in nature (where the environment is rarely constant, both spatially and temporally) (Lande, 1983; Orr and Coyne, 1992; Tenaillon, 2014; Dillon *et al.*, 2016). Our results are in accord with this long-held assumption and lead to the prediction that environmental fluctuations across states that show fitness trade-offs can potentially explain why small populations can be successful in nature. We note that the evolution of ecological specialization may sometimes require thousands of generations (Kassen, 2002, 2014; Cooper, 2014). It would therefore be particularly interesting to study how specialization evolves in populations of different sizes if the environment fluctuates at much smaller timescales (tens of generations). Our results can act as stepping-stones for more complex investigations of the links between population size and trade-offs, particularly in fluctuating environments.

## Acknowledgements

We thank Milind Watve and MS Madhusudhan for their valuable inputs. YDC was supported by a Senior Research Fellowship initially sponsored by IISER Pune and then by Council for Scientific and Industrial Research (CSIR), Govt. of India. SM was supported by an INSPIRE undergraduate fellowship, sponsored by Department of Science and Technology (DST), Govt. of India. This project was supported by an external grant (BT/PR22328/BRB/10/1569/2016) from Department of Biotechnology, Govt. of India, and internal funding from IISER Pune.

## Data archiving

All the data relevant to this study would be made publicly available in the Dryad Digital Repository upon acceptance.

## Supplementary Information

## Appendix S1: Ecological specialization can happen with or without costs of adaptation

The usage of the term ‘trade-off’ itself has been quite inconsistent across evolutionary studies over the last few decades, particularly in the context of ecological specialization (Fry, 1996; Bell and Reboud, 1997; Kassen, 2002; Fry, 2003; Gwynn *et al.*, 2005; Jessup and Bohannan, 2008; Lang *et al.*, 2009; Smith-Tsurkan *et al.*, 2010; Kassen, 2014). On the one hand, the term ‘trade-off’ has been used interchangeably with ‘cost of adaptation’, which represents cases where adaptation to one environment leads to maladaptation in another. Such a notion suggests that maladaptation (decrease in fitness below the ancestral level) is a pre-requisite for trade-offs (Kawecki *et al.*, 1997; Bohannan *et al.*, 2002; Fry, 2003; Smith-Tsurkan *et al.*, 2010). On the other hand, some studies explicitly differentiate between trade-offs and costs of adaptation, stating that trade-offs (and thus specialization) can occur with or without such costs (Bell and Reboud, 1997; Kassen, 2014). Such discrepancy in the usage of the term trade-off can often lead to confusions while comparing the conclusions of different studies. For example, single-generation studies of trade-offs and ecological specialization routinely make conclusions about costs of adaptation while relying almost exclusively on negative correlations in fitness across environments (Joshi and Thompson, 1995; Fry, 2003; Jessup and Bohannan, 2008; Maharjan *et al.*, 2013). However, single generation studies are not only weak in terms of detecting specialization, they can also make misleading claims regarding costs of adaptation (Kassen, 2002; Fry, 2003).

As shown schematically in Fig. 1, negative fitness correlations only imply the intersection of competing reaction norms for fitness across the two environments in question. Such intersection only means that no genotype has the highest fitness in both the environments. More importantly, reaction norms can intersect (and thus fitness correlations can be negative) even if the population ends up adapting to both the environments, albeit to different degrees (Fig. 1a) (Fry, 1996; Kassen, 2014). Thus, although negative correlations always lead to specialization across two environments, they can evolve with or without costs of adaptation (Bell and Reboud, 1997; Kassen, 2002, 2014) (Fig. 1). Thus, ecological specialization can happen even in the absence of costs of adaptation brought about by antagonistic pleiotropy. Indeed, magnitude pleiotropy (cases when the fitness effects of a mutation have the same sign but different magnitudes across environments) can lead to ecological specialization on its own (Remold, 2012). Such magnitude pleiotropy has been a very common observation in recent studies of mutational fitness effects across environments (Schick *et al.*, 2015; Sane *et al.*, 2018).

**Fig. S1.**
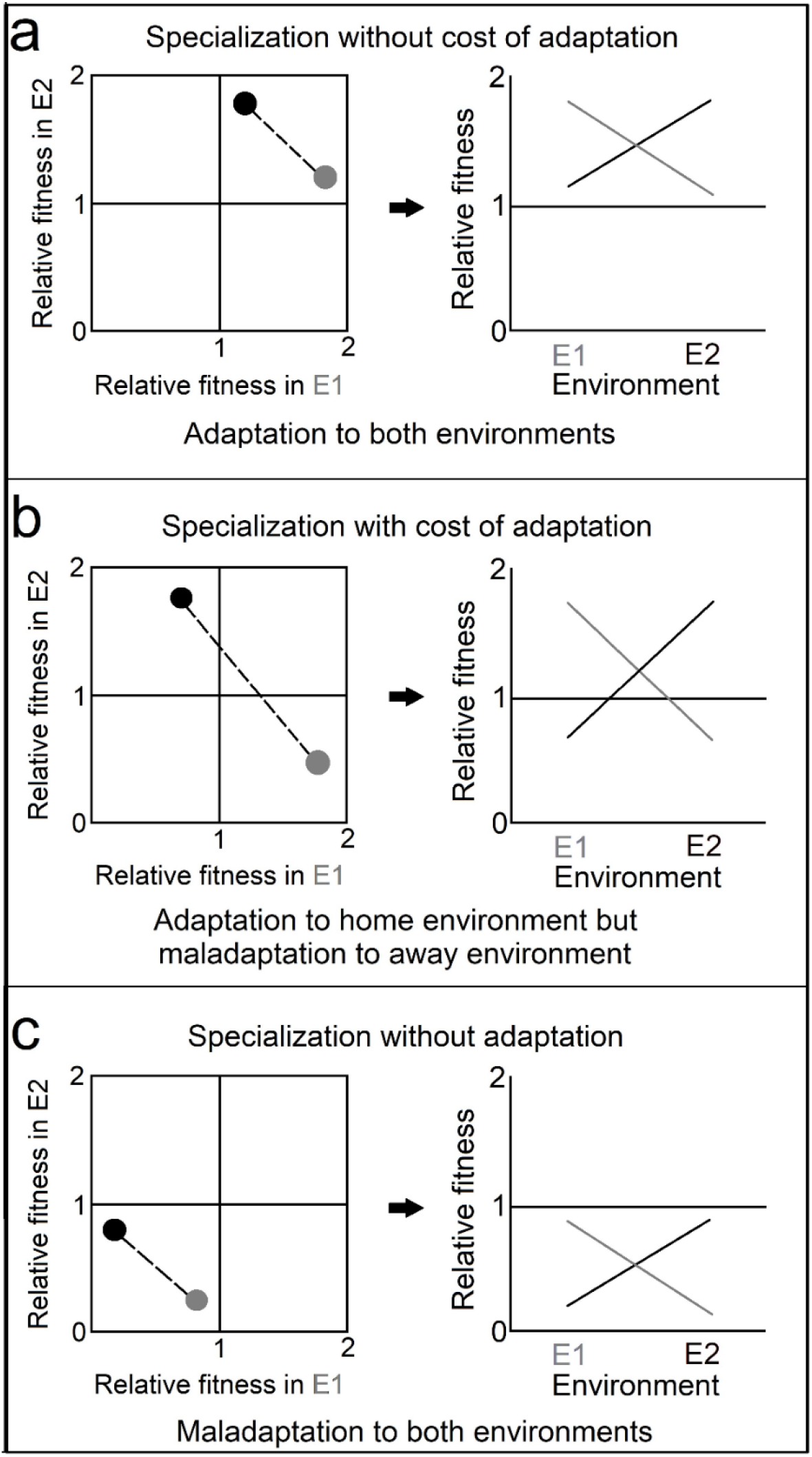
The relationship between ecological specialization and costs of adaptation. Specialization happens when the fittest genotype in one environment is not the fittest genotype in another environment. We consider specialization across two environments (E1 and E2) here. The grey populations evolved only in E1 while the black populations evolved only in E2. The first column across the three panels (a, b, and c) shows a negative fitness correlation in reciprocally selected lines. The second column shows the reaction norms corresponding to the correlation in the first column. Following (Kassen, 2014), the fitness values have been normalized by the ancestral fitness (=1) in each environment. **(a)** Specialization can happen even if reciprocally selected lines end up adapting to both the environments. **(b)** Specialization can happen if reciprocally selected lines adapt to their selective conditions but maladapt to the alternative environment. **(c)** Specialization can happen even if reciprocally selected lines end up maladapting to both the environments.

## Appendix S2: Details of the ancestral strain and media compositions

## Ancestral strain

*Escherichia coli* MG1655 lacY::kan (resistant to kanamycin).

## Composition of the minimal media

Our experiment involved four different M9-based minimal media, each containing one of the following as the only source of carbon:

1. Thymidine
2. Galactose
3. Maltose
4. Sorbitol

1 litre of each minimal medium contained the following:

- 12.8 g Na_2_HPO_4_.7H_2_O
- 3.0 g KH_2_PO_4_
- 0.5 g NaCl
- 1.0 g NH_4_Cl
- 240.6 mg MgSO_4_
- 11.1 mg CaCl_2_
- 4g of the pre-decided carbon source
- 50 mg Kanamycin sulphate

## Appendix S3: Supplementary results

## Single-sample t-test results

**Table S1.** Single-sample t-tests against the ancestral fitness values (N = 6). The fourth column shows Holm-Šidák corrected *P* values. Effect sizes were interpreted as the following: 0.2 < *d* < 0.5 (small effect), 0.5 < *d* < 0.8 (medium effect); *d* > 0.8 (large effect). Effect sizes were not interpreted for cases where Holm-Šidák corrected *P* > 0.05.

| Population type | Assay environment | *P* value | Corrected *P* value | Cohen's *d* (Effect size) |
| --- | --- | --- | --- | --- |
| TL | Thy | $8.100 \times 10^{-7}$ | $3.240 \times 10^{-6}$ | 12.129 (large) |
| TL | Gal | $8.943 \times 10^{-5}$ | $2.683 \times 10^{-4}$ | - 4.670 (large) |
| TL | Mal | $5.553 \times 10^{-4}$ | $5.553 \times 10^{-4}$ | - 3.187 (large) |
| TL | Sor | $1.135 \times 10^{-4}$ | $2.270 \times 10^{-4}$ | - 4.445 (large) |
| TS | Thy | $1.832 \times 10^{-4}$ | $7.327 \times 10^{-4}$ | 4.024 (large) |
| TS | Gal | $6.839 \times 10^{-3}$ | $6.839 \times 10^{-3}$ | - 1.807 (large) |
| TS | Mal | $4.576 \times 10^{-4}$ | $1.372 \times 10^{-3}$ | - 3.318 (large) |
| TS | Sor | $6.190 \times 10^{-4}$ | $1.238 \times 10^{-3}$ | - 3.111 (large) |
| GL | Thy | 0.003 | 0.013 | - 2.158 (large) |
| GL | Gal | 0.016 | 0.049 | 1.448 (large) |
| GL | Mal | 0.633 | 0.633 | - |
| GL | Sor | 0.973 | 0.973 | - |
| GS | Thy | 0.122 | 0.122 | - |
| GS | Gal | 0.617 | 0.617 | - |
| GS | Mal | 0.483 | 0.483 | - |
| GS | Sor | 0.016 | 0.061 | - |

**Table S2.** Single-sample t-tests against the ancestral reaction norm slopes (N = 6). The fourth column shows Holm-Šidák corrected *P* values. Effect sizes were interpreted as the following: 0.2 < *d* < 0.5 (small effect), 0.5 < *d* < 0.8 (medium effect); *d* > 0.8 (large effect). Effect sizes were not interpreted for cases where Holm-Šidák corrected *P* > 0.05.

| Population type | Home-Away pair | *P* value | Corrected *P* value | Cohen's *d* (Effect size) |
| --- | --- | --- | --- | --- |
| TL | T-G | $6.481 \times 10^{-8}$ | $6.481 \times 10^{-8}$ | 20.131 (large) |
| TL | T-M | $2.467 \times 10^{-8}$ | $7.402 \times 10^{-8}$ | 24.427 (large) |
| TL | T-S | $5.935 \times 10^{-8}$ | $1.187 \times 10^{-7}$ | 20.490 (large) |
| TS | T-G | $5.633 \times 10^{-5}$ | $5.633 \times 10^{-5}$ | 5.136 (large) |
| TS | T-M | $5.490 \times 10^{-5}$ | $1.098 \times 10^{-4}$ | 5.163 (large) |
| TS | T-S | $4.714 \times 10^{-5}$ | $1.414 \times 10^{-4}$ | 5.328 (large) |
| GL | G-T | $1.830 \times 10^{-4}$ | $5.488 \times 10^{-4}$ | 4.025 (large) |
| GL | G-M | $2.217 \times 10^{-4}$ | $4.434 \times 10^{-4}$ | 3.866 (large) |
| GL | G-S | $5.910 \times 10^{-3}$ | $5.910 \times 10^{-3}$ | 1.873 (large) |
| GS | G-T | 0.232 | 0.232 | - |
| GS | G-M | 0.504 | 0.504 | - |
| GS | G-S | 0.061 | 0.061 | - |

